# Conserved and non-conserved triggers of 24-nt reproductive phasiRNAs in eudicots

**DOI:** 10.1101/2021.01.20.427321

**Authors:** Suresh Pokhrel, Kun Huang, Blake C. Meyers

## Abstract

Plant small RNAs (sRNAs) play important roles in plant growth and development by modulating expression of genes and transposons. In many flowering plant species, male reproductive organs, the anthers, produce abundant phased small interfering RNAs (phasiRNAs). Two classes of reproductive phasiRNAs are generally known, mostly from monocots: pre-meiotic 21-nt phasiRNAs triggered by miR2118, and meiotic 24-nt phasiRNAs triggered by miR2275. Here, we describe conserved and non-conserved triggers of 24-nt phasiRNAs in several eudicots. We found that the abundant 24-nt phasiRNAs in the basal eudicot columbine are produced by the canonical trigger, miR2275, as well as by other non-conserved triggers, miR482/2118 and aco_cand81. These triggering miRNAs are localized in microspore mother cells (MMC) and tapetal cells of meiotic and post-meiotic stage anthers. Furthermore, we identified a new trigger (miR11308) of 24-nt phasiRNAs and an expanded number of 24-*PHAS* loci in wild strawberry. We validated the presence of miR2275-derived 24-nt phasiRNAs pathway in rose. Finally, we evaluated all the eudicots that have been validated for the presence of 24-nt phasiRNAs as models to study biogenesis and function of 24-nt phasiRNAs and conclude that columbine would be an excellent model because of its extensive number of 24-*PHAS* loci and its diversity of trigger miRNAs.

## Introduction

Plant sRNAs range from 20 to 24 nucleotides (nt) in length and are mainly classified into three groups based on their origin and biogenesis. The groups are (1) microRNAs (miRNAs), (2) heterochromatic small interfering RNAs (hc-siRNAs), and (3) phasiRNAs (Fei et al., 2013). Plant sRNA biogenesis requires RNA-DEPENDENT RNA POLYMERASE (RDR), DICER-LIKE (DCL), and Argonaute (AGO), proteins, plus other accessory proteins. All types of sRNAs associate with AGO proteins to initiate both transcriptional and post-transcriptional silencing via formation of RNA-induced silencing complexes. Among these sRNA classes, phasiRNAs are generated by the action of 22-nt miRNA triggers on messenger RNA (mRNA) targets derived from RNA Polymerase II; the target is subsequently processed by RDR6 and DCL proteins into 21- or 24-nt siRNAs (Fei et al., 2013).

Reproductive 24-nt phasiRNAs were reported mainly in the male organs of grasses (Poaceae) (Zhai et al., 2015b; Song et al., 2012; Johnson et al., 2009; Patel et al., 2020), and more recently in six non-grass monocots (Patel et al., 2020) and eight eudicots (Xia et al., 2019). This size class is abundant in the meiotic stage of anthers and are mainly triggered by miR2275, although there are cases of trigger-less 24-nt phasiRNA-producing loci (aka “24-*PHAS*” loci) in tomato and petunia (Xia et al., 2019), and in monocots outside the Poaceae (Kakrana et al., 2018; Patel et al., 2020). The spatiotemporal pattern of 24-nt phasiRNAs in male reproductive tissues has been well-described in maize, rice, and litchi (Zhai et al., 2015b; Song et al., 2012; Xia et al., 2019). The loci that generate the 24-nt phasiRNAs, known as 24-*PHAS* loci, correspond to unique or low-copy genomic regions which makes 24-nt phasiRNAs different from 24-nt heterochromatic siRNAs (hc-siRNAs) that function to suppress transposons (Zhai et al., 2015a). These reproductive 24-nt phasiRNAs lack obvious targets but play a role in male fertility, as their absence due to a knockout of *DCL5* in maize produces temperature-sensitive genic male sterility (Teng et al., 2020). The 24-nt reproductive phasiRNAs may be comparable or analogous to animal piRNAs (PIWI-interacting RNAs) which also play a role in male fertility (Kotov et al., 2019). The role of animal piRNAs is not entirely clear and is an active topic of investigation by numerous labs; however, at least some piRNAs function as a defense mechanism to protect genetic information in germ cells from transposons (Kotov et al., 2019).

Recently, we identified 24-nt phasiRNAs in seven eudicots, demonstrating the broad presence of the miR2275/24-nt phasiRNA pathway in angiosperms (Xia et al., 2019). The general lack of information on this pathway in eudicots motivated us study more species and in more detail. Here, we report the existence of the 24-nt phasiRNA pathway in the basal eudicot columbine (*Aquilegia coerulea*), expand the number of known 24-*PHAS* loci in wild strawberry (*Fragaria vesca*), and validate the pathway in rose (*Rosa chinensis*). We identify triggers of 24-nt phasiRNAs other than miR2275, in columbine and wild strawberry. Finally, we evaluated the eudicot species validated for the presence of the 24-PHASiRNA pathway by potential characteristics, with the aim to identify a model species in which to study biogenesis and function of this pathway; we conclude that columbine would be an excellent model for these studies.

## Results

### Conserved triggers initiate 24-nt phasiRNAs in columbine anthers

miR2275 is mostly conserved in angiosperms and is the only trigger of 24-nt phasiRNAs known to date (Xia et al., 2019). Using this as a marker of the 24-nt phasiRNA pathway, we searched for the presence of miR2275 in the genome of columbine and identified 11 clusters of miR2275 precursors that yield 19 mature duplexes with 12 unique, expressed, miR2275 sequences (Figure 1A, Supplemental Table 1). Out of the 11 clusters, six are polycistronic precursors; Figure 1B shows an example of a polycistronic cluster. To study 24-nt phasiRNA expression in columbine, we analyzed sRNAs data derived from vegetative tissues (leaf and root) and reproductive tissues (four unopened bud stages; Pokhrel et al., 2020). We found that mature miR2275 sequences highly accumulate in bud stages two and three (Figure 1C) which represent meiotic/post-meiotic stages of anther development. We identified an extensive number (642) of 24-*PHAS* loci with miR2275 target sites as the only apparently conserved sequence among the precursors (Supplemental Figure 1A, Supplemental Table 2). The 24-nt phasiRNAs spawned from these precursors demonstrate similar patterns of abundance as their triggers, suggesting a tight temporal regulation as in grasses and litchi (Figure 1D; Zhai et al., 2015b; Xia et al., 2019). The miR2275-triggered 24-*PHAS* loci are highly phased and the first 24-nt phasiRNA originates from cleavage site indicated by the red arrow in an example locus as shown (Supplemental Figure 1B).

**Table 1.** Eudicots confirmed for the presence of 24-nt phasiRNA pathway.

| Species – common name or shorthand | Scientific name | Family | Genome sequenced? | Genome size | Plant life cycle | No of 24-nt PHAS loci | miRNA triggers of reproductive phasiRNAs | No. of miR2275 duplexes | Genomic copies of *DCL3* | References |
| --- | --- | --- | --- | --- | --- | --- | --- | --- | --- | --- |
| Columbine | *Aquilegia coerulea* | Ranunculaceae | Yes | ~300 Mb | Perennial | 646 | miR2275, aco_cand81, miR2118/482 | 15 | 2 | This study |
| Grape | *Vitis vinifera* | Vitaceae | Yes | ~487 Mb | Perennial | 42 | miR2275 | 10 | 1 | Xia *et al.*, 2019 |
| Rose | *Rosa chinensis* | Rosaceae | Yes | ~503 Mb | Perennial | 112 | miR2275 | 10 | 2 | This study |
| Strawberry | *Fragaria vesca* | Rosaceae | Yes | ~240 Mb | Annual | 335 | miR2275, miR11308, miR2118/482 | 17 | 2 | Xia *et al.*, 2019; this study |
| Citrus | *Citrus sinensis* | Rutaceae | Yes | ~367 Mb | Perennial | 32 | miR2275 | 10 | 2 | Xia *et al.*, 2019 |
| Litchi | *Litchi chinensis* | Sapindaceae | No | ~550-620 Mb | Perennial | 113 | miR2275 | 6 | 2 | Xia *et al.*, 2019 |
| Cotton | *Gossypium hirsutum* | Malvaceae | Yes | ~2,295 Mb | Annual | 3 | miR2275 | 1 | 2 | Xia *et al.*, 2019 |
| Tomato | *Solanum lyopersicum* | Solanaceae | Yes | ~950 Mb | Annual | 137 | None | None | 1 | Xia *et al.*, 2019 |
| Petunia_ax | *Petunia axillaris* | Solanaceae | Yes | ~181 Mb | Perennial | 317 | None | None | 1 | Xia *et al.*, 2019 |
| Petunia_inf | *Petunia inflata* | Solanaceae | Yes | ~197 Mb | Perennial | 366 | None | None | 1 | Xia *et al.*, 2019 |

**Figure 1.**
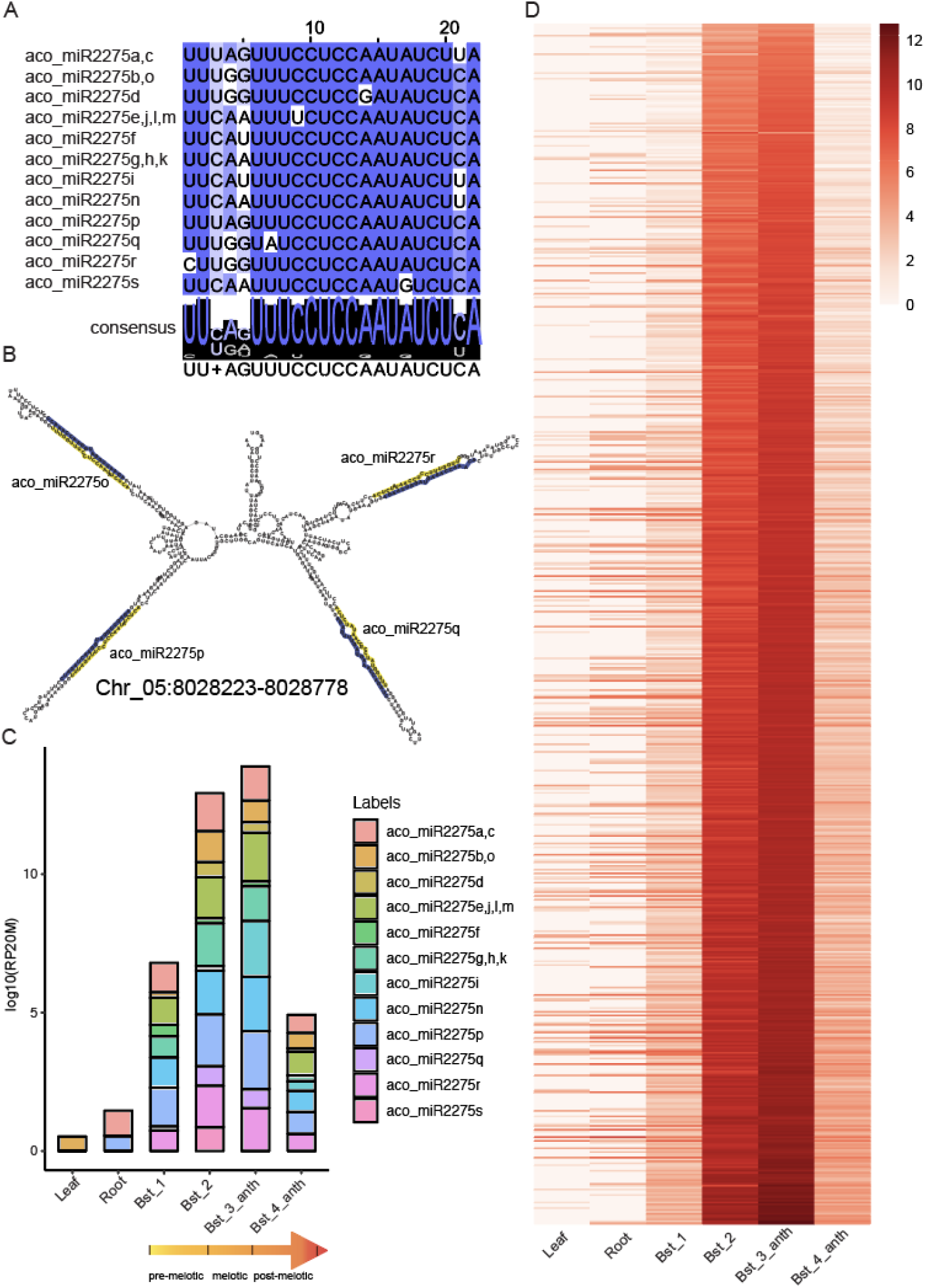
Reproductive 24-nt phasiRNAs triggered by miR2275 are abundant in columbine. **A.** Alignment of miR2275 family members in columbine. The intensity of blue color denotes the level of conservation, with the darker color indicating a more highly conserved sequence; the consensus sequence is shown with a “sequence logo”. **B.** An example polycistronic cluster of miR2275 members in columbine. The mature and star sequences are marked in yellow and blue respectively. Coordinates of the precursor are indicated below the sequences. **C.** Abundance of miR2275 members in log10(RP20M) in different tissues of columbine. The yellow gradient arrow at the bottom shows the progression of anther development along different stages. **D.** Expression of miR2275-triggered 24-nt reproductive phasiRNAs in vegetative and reproductive tissues. The key at right indicates the abundance in units of log2(RP20M). Bst indicates bud stage, anth indicates anther. RP20M: reads per twenty million mapped reads (normalized abundance).

### Non-conserved triggers initiate 24-nt phasiRNAs in columbine anthers

During our analysis of possible triggers of 24-*PHAS* loci, we unexpectedly found that both (i) a previously-unannotated, apparently lineage-specific miRNA (aco_cand81; Supplemental Figure 2A) and (ii) miR2118 family members (Supplemental Figure 2B) triggered production of 24-nt phasiRNAs from 15 and 22 loci, respectively (Figure 2A, 2B). The expression profile of the 24-nt phasiRNAs generated via the non-conserved triggers are similar to those derived from the conserved, canonical trigger, miR2275 (Figure 1D, 2A, 2B). Ten out of 15 24-*PHAS* loci targeted by miRNA aco_cand81 are derived from a cluster spanning 32.2 kb on chromosome 4 (Supplemental Figure 2C, Supplemental Table 2), suggesting that these 24-nt phasiRNAs were selectively retained in the genome as the loci don’t resemble the product of a recent tandem duplication -- a whole precursor alignment of these loci shows no conservation at the nucleotide level, except for the target site. That is, the only conserved sequence between these 15 24-*PHAS* loci is the target site of aco_cand81 (Figure 2C). Similarly, the 22 24-*PHAS* loci triggered by miR2118 also showed conservation of only the target site (Figure 2D, Supplemental Table 2). In the genome browser, it is apparent that the first 24-nt phasiRNAs are generated from the cleavage site of aco_can81 and miR2118 (Figure 2E and 2F), respectively. Interestingly, members of both of these miRNA families also trigger production of 21-nt reproductive phasiRNAs in columbine (Pokhrel et al., 2020).

**Figure 2.**
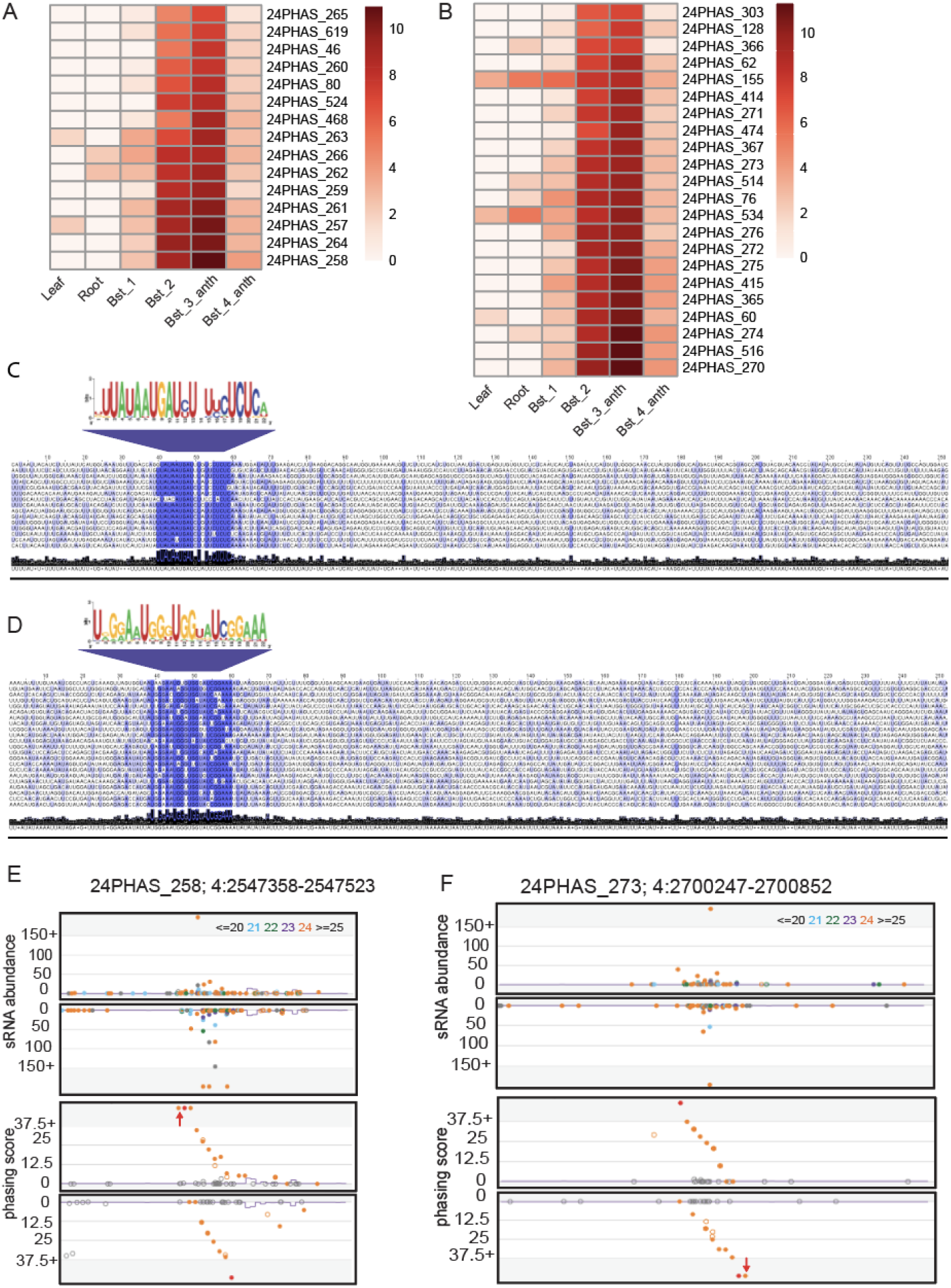
Non conserved triggers generated 24-nt phasiRNAs in columbine. **A.** Expression of aco_cand81-triggered 24-nt reproductive phasiRNAs in vegetative and reproductive tissues. The key at right indicates the abundance in units of log2(RP20M). Bst indicates bud stage, anth indicates anther. **B.** Same as A for miR2118/482 derived 24-*PHAS* loci. **C.** Above: Consensus sequence of target sites of aco_cand81, derived from 15 24-*PHAS* loci. Below: Nucleotide sequence alignment of 15 24-*PHAS* loci with sequence similarity denoted by the intensity of the blue color indicating that the target site is the only conserved region for all precursors. **D.** Same as C for miR2118, derived from 22 24-*PHAS* loci. **E.** Top: Abundance (in RP10M) of small RNAs in both strands of an example locus padded with 1000 base pairs, each side. sRNA sizes are denoted by colors, as indicated at top. Bottom: Phasing score of the same locus; the red dot indicates the highest phased sRNA position. The red arrow denotes cleavage site of aco_cand81. **F.** Same as E for an example 24-*PHAS* locus triggered by miR2118 members.

In the genome of columbine, 24-*PHAS* loci are distributed across the arms of all chromosomes, while one arm of chromosome 4 has a particularly high concentration of loci (313 loci out of 679 total loci; outer circle, Figure 3A). We examined the strand-specificity of 24-*PHAS* precursors (inferring the Pol II-derived strand from the target site match) within the chromosome 4 clusters. We found that a conservation of strandedness within a cluster, meaning that all the precursors are encoded on the same chromosomal strand; however, there is again no conservation of nucleotide sequence other than the target site of the triggering miRNAs, even within these tandemly arranged sequences. We hypothesized that there was a duplication of 24-*PHAS* loci that caused the arrangement (Supplemental Figure 1A; Figure 3B). The duplications must have occurred long enough ago to allow nearly complete sequence divergence; a dot plot of 24-*PHAS* loci from chromosome 4 demonstrates very few similarities among these loci (Supplemental Figure 2D). Chromosome 4 of Aquilegia has previously been shown to have a “distinct evolutionary path from the rest of the genome” (Aköz and Nordborg, 2019), and containing many duplications consistent with the arrangement of the 24-*PHAS* loci. In contrast, the conserved and non-conserved triggers of these phasiRNAs are broadly distributed, on all chromosomes except 2 and 7 (inner circle, Figure 3A).

**Figure 3.**
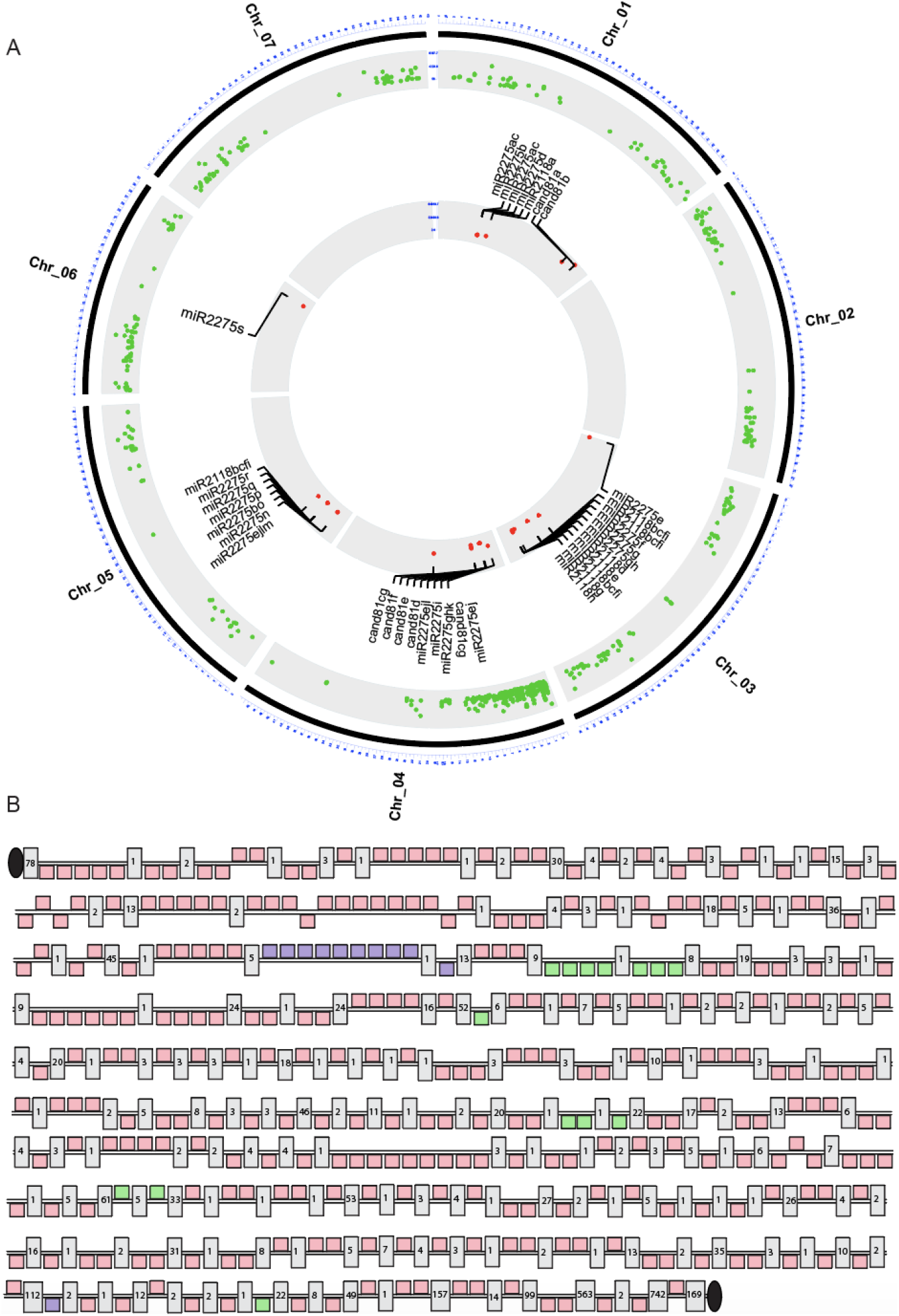
Genome-wide distribution of all triggers and 24-*PHAS* loci in columbine. **A.** Chromosomal positions of miR2275, miR2118/482, and aco_cand81 family members (inner circle, red dots) and their 24-*PHAS* loci (outer circle, green dots). The outermost circle represents chromosomes from 1 to 7. **B.** Chromosomal positions and strand-specificity of 24-*PHAS* loci on chromosome 4 interspersed with genes. The black line represents chromosome 4, and the square boxes above and below the line indicate plus and minus strand-specificity, respectively. The light pink, light purple, and light green boxes represent the 24-*PHAS* loci triggered by miR2275, aco_cand81, and miR2118/482 family members, respectively. Grey rectangle boxes represent annotated protein-coding genes that intersperse the *PHAS* clusters, with the total number of interspersed genes indicated in each box. The black ovals represent “telomeric end” for chromosome 4.

We next profiled the single nucleotide compositions of 24-nt phasiRNAs from columbine, analyzed in three different groups based on their triggers, and compared these to hc-siRNAs. This approach can identify seemingly minor but statistically significant differences in sRNA populations (Patel et al., 2018a). We computed the frequency of each nucleotide (A, C, G and U) at each position (Supplemental Figure 3) for the 24-nt phasiRNAs and hc-siRNAs. We observed position-specific biases, indicating different nucleotide composition, in these two different classes of siRNAs (Supplemental Figure 3A and B). Moreover, we found that the nucleotide composition of 24-nt phasiRNAs derived from different triggers was the same (Supplemental Figure 3C), consistent with shared origins and derivations.

### Spatiotemporal localization of triggers and 24-nt phasiRNAs in columbine

We next examined the spatiotemporal localization of all three reproductive phasiRNA triggers in pre-meiotic, meiotic, and post-meiotic stages of columbine anther development. The non-conserved triggers aco_can81 and miR2118 were enriched compared to miR2275, in all stages. The triggers localized to all cell layers and were enriched in the microspore mother cells (MMC) of pre-meiotic stage or tapetal cells of meiotic/post-meiotic stages, suggesting 24-nt phasiRNAs originate in those cell layers (Figure 4A). Because the imaging method we utilized is a single-molecule, quantitative method, we also quantified the copy number of each phasiRNA trigger in MMC, tapetal cells, and other cell layers. Overall, miR2275 was enriched in the post-meiotic stage compared to other stages, possibly reflecting additional roles it may play in this stage, while miR2118 and aco_cand81 were enriched in meiotic/post-meiotic stages, mirroring the abundance patterns of the 24-nt phasiRNAs they generate (Figure 4B; Figure 2A and 2B). The spatiotemporal pattern of triggering molecules in columbine is consistent with that of litchi and maize (Xia et al., 2019; Zhai et al., 2015b), consistent with a conserved and possibly important regulatory role of these RNAs in male gamete formation.

**Figure 4.**
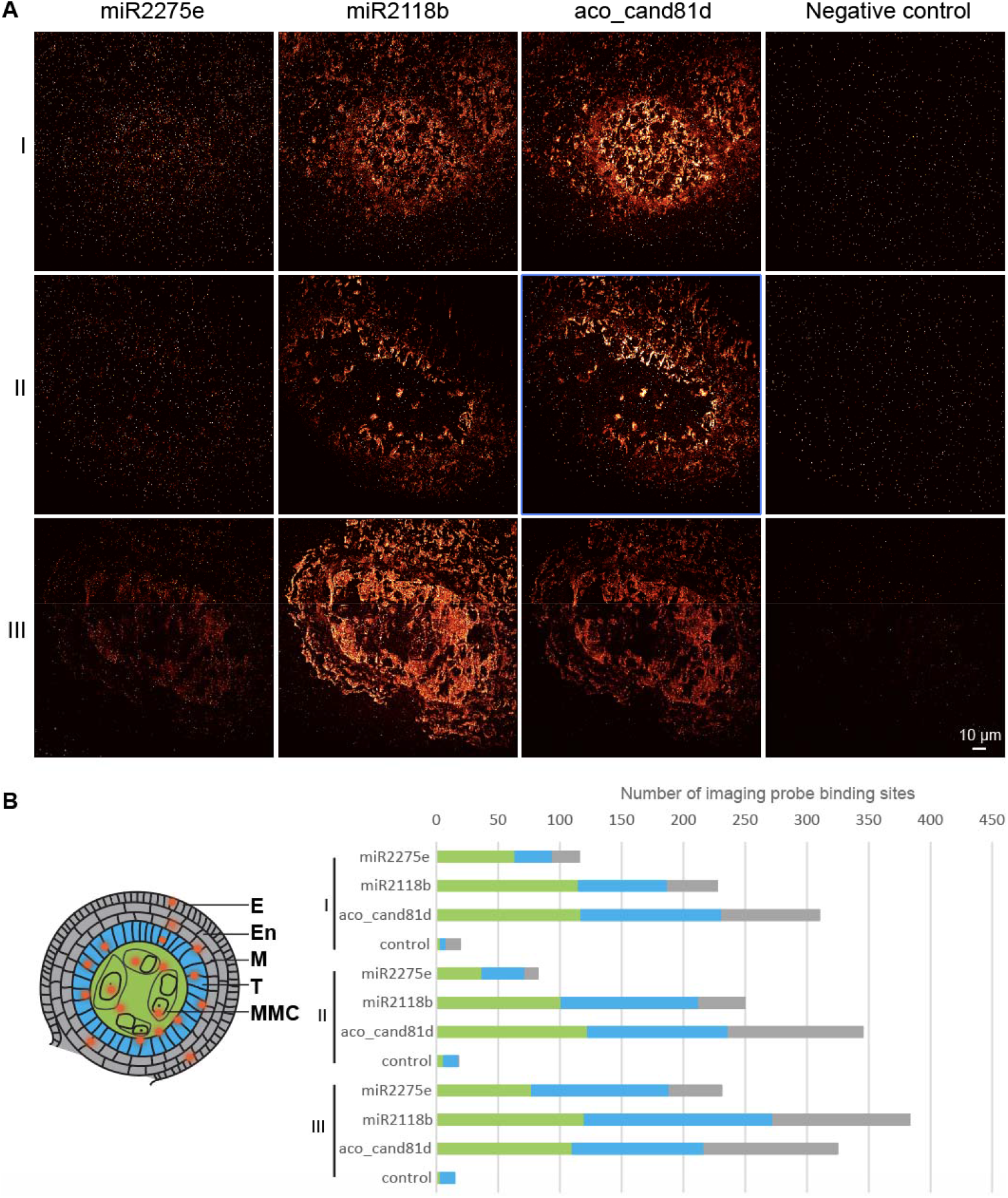
Trigger miRNAs are enriched in microspore mother cells and tapetal cells of anthers in columbine. **A.** sRNA-PAINT of all the triggers in pre-meiotic, meiotic and post-meiotic anther stages in columbine. I, II, and III indicate pre-meiotic, meiotic, and post meiotic stages. In each stage, an image of an anther lobe was captured, with the different cell layers following the pattern shown in the schematic diagram of B. White dots in the negative control images were gold fiducials that are pre-coated on the coverslips used for focal panel stabilization during imaging process. Imager buffer with no imager strands was used as negative controls on the same hybridized slides shown for each stage. Scale bar = 10 ¼m for all images. **B.** Left: schematic diagram of the sRNA-PAINT method applied in a lobe of an anther, showing different cell layers, labeled as follows: E: epidermis, En: endodermis, M: middle layer, T: tapetum, MMC: microspore mother cell. The red dots indicate the sRNAs hybridized to different cell layers. Right: Number of imaging probe binding sites of each trigger in MMC (green), tapetal cells (blue), and other cell type (grey) in pre-meiotic, meiotic, and post-meiotic stages of anther development. The number of binding sites was calculated as an average from ten measurements in a cell (30 pixels) from different cell layers in each sample.

### Conserved and non-conserved triggers of 24-nt phasiRNAs in the Rosaceae and Magnoliids

Recently, we described the presence of a miR2275-triggered, 24-nt phasiRNA pathway in wild strawberry (Xia et al., 2019) with the highest known number (which was 221) of 24-*PHAS* loci in eudicots. By using two types of phasiRNA-identifying software, coupled with extensive experimental data (Xia et al., 2019), we aimed to saturate the identification of 24-*PHAS* loci in wild strawberry. The result was that this analysis expanded the set of known miR2275-triggered 24-*PHAS* loci to 326 in total (Supplemental Figure 4A, Supplemental Table 3). These phasiRNAs were expressed mostly in anther stages 7 to 9, corresponding to the meiotic stages of anther development. The expression pattern of miR2275 variants also followed a similar pattern (Supplemental Figure 4B, 4C). Then, we sought to determine if any additional triggers were previously missed, as prior work focused only on miR2275; we found that miR11308 generated 24-nt phasiRNAs from six 24-*PHAS* loci, while miR482/2118 family members targeted four 24-*PHAS* loci (Figure 5A, Supplemental Table 3). The expression pattern of 24-nt phasiRNAs generated via these non-conserved triggers was similar to that of the canonical trigger, miR2275 (Figure 5A; Supplemental Figure 4A). The target site of miR11308 was the only conserved region in six 24-*PHAS* loci (Figure 5B). These loci mostly originated from arms of all chromosomes and their triggers are encoded by loci found on chromosomes 1, 4, 5 and 6 (Figure 5C).

**Figure 5.**
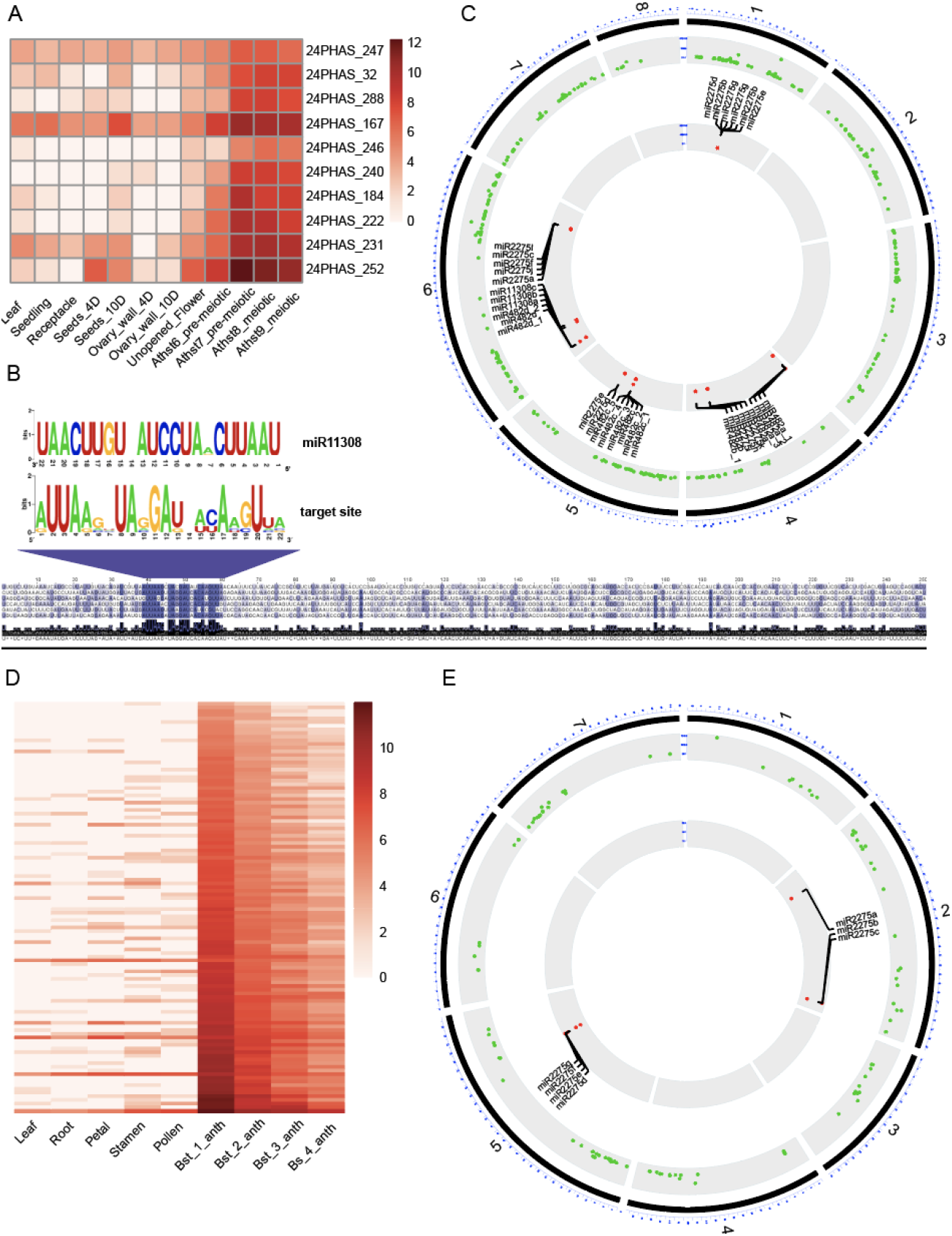
Reproductive 24-nt phasiRNAs and their triggers in the Rosaceae. **A.** Expression of 24-nt reproductive phasiRNAs derived by the action of non-conserved triggers, in different tissues and anther development stages in wild strawberry; Athst indicates anther stage. The key at right indicates the abundance in units of log2(RP20M). **B.** Above: Alignment of sequence logos of miR11308 and target site of miR11308 for six 24-*PHAS* loci. Below: Nucleotide sequence alignment of six 24-*PHAS* loci with sequence similarity denoted by the intensity of the blue color showing that the target site is the only conserved region for all the precursors. **C.** Genome-wide distribution of miR2275, miR11308 and miR2118/482 family members (inner circle, red dots) and their 24-*PHAS* loci (outer circle, green dots). The outermost circle represents chromosomes from 1 to 7 as numbered, plus chromosome “8” which represents an amalgam of the unassembled regions. **D.** Accumulation of 24-nt reproductive phasiRNAs derived by the action of miR2275, in different tissues and anther development stages in rose. The key at right indicates the abundance in units of log2(RP20M); Bst indicates bud stage, and anth indicates anther. **E.** Genome-wide distribution of miR2275 family members (inner circle, red dots) and their 24-*PHAS* loci (outer circle, green dots). The outermost circle represents chromosomes from 1 to 7, as numbered.

To investigate the conservation of this pathway, we analyzed sRNA data of vegetative tissues, and anthers from four unopened bud stages of rose (Guo et al., 2019; Pokhrel et al., 2020). We found 112 24-*PHAS* loci triggered by miR2275 variants, accumulating to the greatest abundance at bud stage one (Figure 5D, Supplemental Table 4). Genomic searches for the presence of miR2275 in rose identified three clusters of precursors, two of which were apparently polycistronic; an example locus is shown in Supplemental Figure 5A (see also Supplemental Table 5). We observed the expression of seven mature sequences of miR2275, and their abundance patterns were similar to that of 24-nt phasiRNAs (Supplemental Figure 5B and 5C). We found no evidence for the non-conserved triggers of strawberry (i.e. miR2118/42 and miR11308) targeting 24-*PHAS* loci in rose, perhaps due to species-specific variation, or possibly due to inadequate sampling of anther stages. Similar to wild strawberry, the *PHAS* loci are distributed across arms of all chromosomes, while the miRNA triggers originated from chromosomes 2 and 5 (Figure 5E).

We previously characterized the presence of miR2275 loci in 209 genomes of flowering plants spanning different clades of eudicots and monocots (Xia et al., 2019); however, that analysis excluded the magnoliids as there were no sequenced genomes at that time. Therefore, we investigated a recently published sequence genome of a magnoliid species, *Magnolia biondii*, (Dong et al., 2020). We found nine unique mature miR2275 sequences derived from 11 precursors (Supplemental Figure 6A), three of which appear to be polycistronic precursors, an example locus is shown in Supplemental Figure 6B (see also Supplemental Table 5).

**Figure 6.**
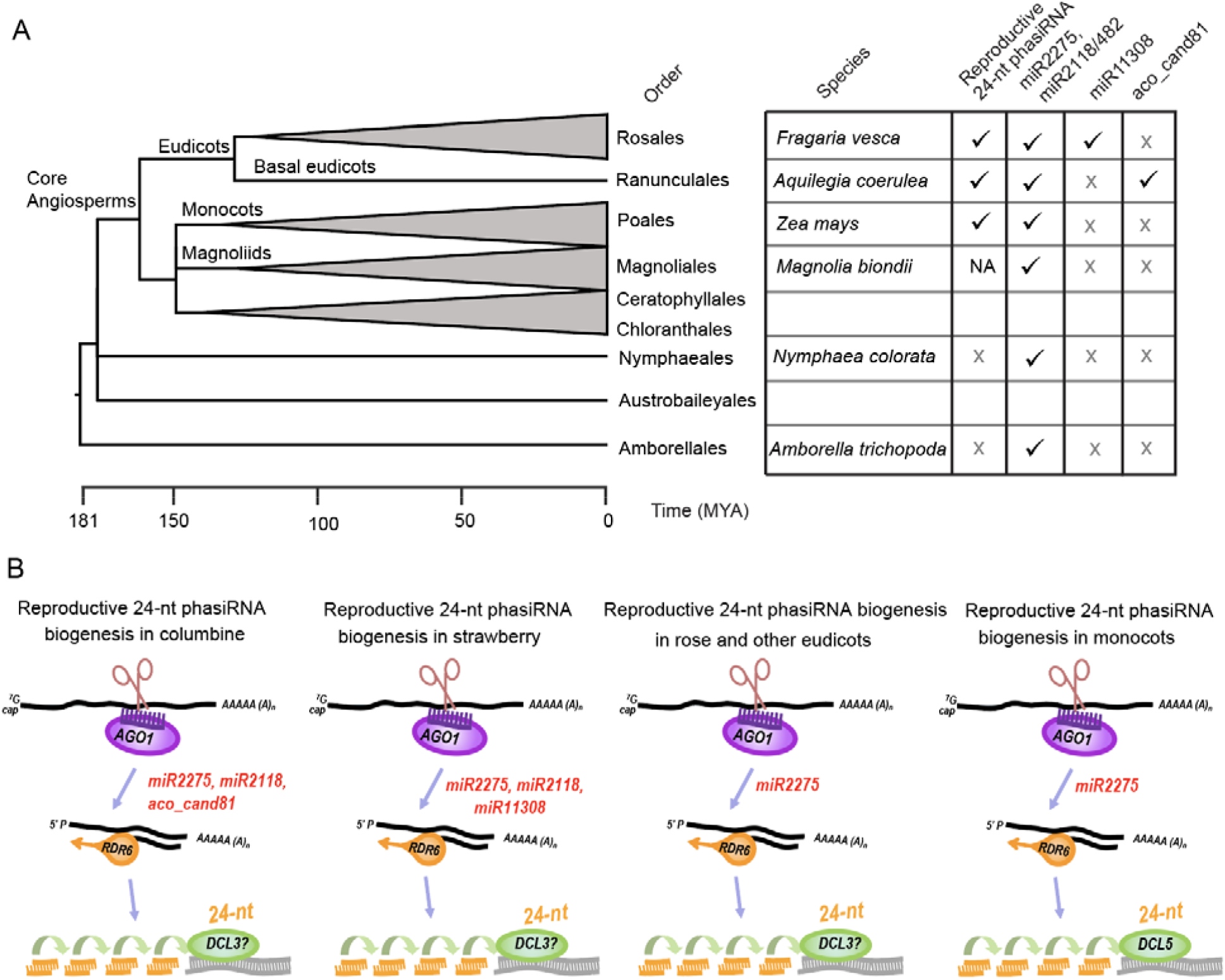
Conservation of reproductive 24-nt phasiRNAs in angiosperms. **A.** Phylogeny showing the representative orders and species of angiosperms. Only one order and one species from each monophyletic clade of eudicots, monocots, and magnoliids was chosen. Phylogenetic tree was modified from the timetree database. MYA = million years ago. In the table at right, a check mark indicates presence and X indicates absence; NA indicates ‘not analyzed’. Species name from basal angiosperm orders left empty because they lack sequenced genomes for analysis. **B.** Model for generation of 24-nt reproductive phasiRNAs in angiosperms in terms of trigger diversity and Dicer processing. There are at least three miRNA family members involved in generating 24-nt phasiRNAs in columbine and strawberry. In rose, other eudicots, and monocots, miR2275 is the only trigger. DCL5 is monocot specific, so DCL3 may function in dicing 24-nt phasiRNAs in eudicots.

### Evaluation of eudicot species as models to study 24-nt phasiRNAs

In total, from this and prior work, the production of 24-nt phasiRNAs was validated in reproductive tissues of ten eudicots (Table 1). To enable further insights into potential mechanistic roles of these phasiRNAs in reproduction, we sought to identify potential model species for future work. Thus, we compared different characteristics such as genome size, plant life cycle, the number of 24-*PHAS* loci and miRNA triggers, and the number of genomic DCL3 copies of the validated eudicot species. Of these species, transformation by tissue culture has been widely reported for all but columbine. Based on this comparison, we conclude that columbine would be an excellent model system due to its phylogenetic position among flowering plants as a basal eudicot, its diversity of trigger miRNAs, and its extensive number of 24-*PHAS* loci. Although it lacks an efficient transformation system, the recent use of developmental genes encoding GROWTH-REGULATING FACTORS such as *AtGRF5* or *TaGRF4* may help to generate genetically modified plants in recalcitrant eudicots such as columbine (Kong et al., 2020; Debernardi et al., 2020). From the perspective of having an already established transformation system and its ease of propagation, wild strawberry may be a good model system, as it also has a diversity of triggers and extensive 24-*PHAS* loci, though fewer compared to columbine.

## Discussion

We showed that the 24-nt reproductive phasiRNA pathway is conserved in a basal eudicot, columbine. Recently, we hypothesized that this pathway emerged coincident with angiosperms with selective maintenance or loss in some flowering plant lineages (Xia et al., 2019). This was based in part on the presence/absence of a canonical pathway trigger (miR2275). The basal angiosperms *Amborella trichopoda and Nymphaea colorata* are the only species outside of eudicots and monocots that have the trigger encoded in their genomes but lack *bona fide 24-PHAS* loci (Xia et al., 2019; Patel et al., 2020). Our confirmation of the presence of this pathway in columbine, an early diverging lineage of the eudicots, provides strong support for the emergence of the pathway prior to the emergence of the core eudicots. Furthermore, in addition to wild strawberry, we validated the presence of this pathway in another *Rosaceae* family member, rose, providing additional evidence for widespread conservation of this pathway. We also identified the presence of miR2275 in *Magnolia biondii*, a magnoliid, suggesting an origin for this pathway of at least 150 MYA in core angiosperms (Figure 6A).

Surprisingly, we found lineage-specific triggers that generate 24-nt phasiRNAs in both columbine and wild strawberry, distinct from the known, canonical trigger, miR2275. miR2275 is deeply conserved in angiosperms, including the characteristic of its polycistronic precursors, and the only known role of miR2275 is the generation of reproductive 24-nt phasiRNAs (Xia et al., 2019). Interestingly, a well-studied miRNA, miR2118/482, known for triggering 21-nt phasiRNAs from both protein-coding and non-coding loci in eudicots and monocots (Zhai et al., 2015b; Bélanger et al., 2020; Pokhrel et al., 2020), has a function in generating 24-nt phasiRNAs in columbine and wild strawberry, adding to the data from Solanaceous species (Xia et al., 2019), barley and wheat (Bélanger et al., 2020) that there are multiple mechanisms for activation of the 24-*PHAS* pathway independent of miR2275. However, there were few 24-*PHAS* loci triggered by miR2118/482 compared to the total number 24-*PHAS* loci in each species. miRNAs that may be lineage-specific, aco_cand81 in columbine and miR11308 in wild strawberry, trigger both 24- and 21-nt reproductive phasiRNAs (Pokhrel et al., 2020). The 21-nt phasiRNAs are produced by the action of DCL4 while DCL3 and DCL5 are responsible for 24-nt phasiRNA production in eudicots and monocots, respectively. DCL5 resulted from a monocot-specific duplication of DCL3 and is specialized for the production of reproductive 24-nt phasiRNAs in monocots (Teng et al., 2020; Patel et al., 2020). Although DCL3 is single copy in the genome of many eudicots, some eudicot lineages that have a duplicated copy of DCL3 are also predicted or validated for the presence of 24-nt phasiRNAs, perhaps consistent with a role for the duplicated DCL3 copy in the biogenesis of reproductive 24-nt phasiRNAs (Pokhrel et al., 2020; Xia et al., 2019). An interesting question for future work will be how different Dicer proteins are recruited by the action of these different trigger miRNAs. Another interesting topic for future work is the role of the polycistronic nature of the precursors observed for miRNA triggers of both 21 and 24-nt reproductive phasiRNAs -- the conserved nature of this characteristic suggests a functional role in reproductive phasiRNA biogenesis. Yet another topic for investigation builds on our observations about the unusual concentration of reproductive PHAS loci on chromosome 4 of columbine. Our data show that the 24-*PHAS* loci have undergone extensive tandem and segmental duplications on chromosome 4. Since the 24-*PHAS* loci are not repetitive DNA, and there are relatively few other genes interspersed among the large number of 24-*PHAS* loci, it is entirely possible that this unusual chromosomal structure and distinct evolutionary path of chromosome 4 (Aköz and Nordborg, 2019) has been driven by and is the consequence of expansion in the 24-*PHAS* loci.

We conclude with a model describing how different triggers produce 24-nt phasiRNAs in different species of angiosperms (Figure 6B); this includes the hypothesis that the activity of the lineage-specific miRNA triggers neofunctionalized to allow targeting precursors of both 21- and 24-nt reproductive phasiRNAs.

Taken together, these data indicate that conservation of 24-nt reproductive phasiRNAs in eudicots is widespread, that they originated coincident with the evolutionary emergence of angiosperms, and that diversification in their mode of biogenesis has occurred. Although the functions of these 24-mers are still unclear, as they lack obvious target genes (Patel et al., 2018b), mutations in a pathway gene, *DCL5*, in maize produced temperature sensitive male sterility, consistent with important regulatory roles for these sRNAs in reproductive development. Future studies are needed to advance our mechanistic understanding of these sRNAs. Based on its phylogenic position as an early-diverged eudicot, the diversity of the miRNA triggers, and the extensive number of 24-*PHAS* loci, we propose that columbine could serve as an ideal model species with which to study these phasiRNAs in eudicots; it lacks only a facile transformation system.

## Methods

### Small RNA datasets

The sRNA-seq data used in this study were published previously (Pokhrel et al., 2020; Xia et al., 2019; Guo et al., 2019; Xia et al., 2015). Briefly, in those studies, sRNA-seq was carried out on vegetative tissues, unopened flower buds/anthers of columbine, rose, and wild strawberry.

### Data analysis

sRNA-seq data analysis was performed as described (Pokhrel et al., 2020). Briefly, an in-house preprocessing pipeline (Mathioni et al., 2017) was used to process the raw reads. Saturating annotation of *PHAS* loci in all of the species was attempted by the combined use of two software *PHASIS* (Kakrana et al., 2017) and ShortStack (Johnson et al., 2016). The *PHAS* loci from these two different software packages were merged using BEDTools (Quinlan and Hall, 2010). Target prediction was carried out using sPARTA (Kakrana et al., 2014) and the small RNA abundances and phasing scores were viewed on the Meyers lab genome browser (Nakano et al., 2020) to filter out the false-positive *PHAS* loci. Only *PHAS* loci having conserved target sites for miRNA triggers were annotated as valid. miR2275 variants from columbine were aligned with genome of *Magnolia biondii* using bowtie (Langmead et al., 2009) with default parameters except –v 2. The resulting mapped positions were padded by 100 bp on each side, and the sequences were extracted using BEDTools (Quinlan and Hall, 2010); RNAfold (Hofacker, 2003) was used to assess the folding of the sequences to identify the most likely precursors of miR2275 encoded in the genome (Supplemental Table 5).

### Data visualization

Multiple sequence alignment of strand-specific sequences from PHAS loci was performed using MUSCLE (Edgar, 2004) with default parameters. The resulting alignment was viewed using Jalview (Waterhouse et al., 2009). Sequences extracted from the alignment were used to construct a dot plot of 24-*PHAS* loci from chromosome 4 in columbine using D-Genies (Cabanettes and Klopp, 2018). Position-specific nucleotide biases were calculated using PWM_StatisicalModel (Patel et al., 2018b). Circular plots were made using OmicCircos (Hu et al., 2014) for the chromosomal distributions. In R, pheatmap (https://rdrr.io/cran/pheatmap/) and ggpubr (https://github.com/kassambara/ggpubr) were used to draw heatmaps and box-plot. The phylogenetic tree was derived from the TimeTree database (Kumar et al., 2017) and modified using FigTree (http://tree.bio.ed.ac.uk/software/figtree/).

### sRNA imaging

miRNA triggers aco_cand81d, miR2275e, and miR2118b were detected in anther sections using sRNA-Exchange-PAINT as described in the published protocol (Huang et al., 2020). Probes for miRNAs were designed using the VARNISH website (https://wasabi.ddpsc.org/~apps/varnish/). Probes and corresponding imager strands with AlexFluor647 were ordered from IDT (Integrated DNA Technologies, Inc.; sequences in Supplemental Table 6). Paraffin-embedded samples of carefully-staged flower buds were sectioned and dried on Wide Spectral Band 600+/−100 nm Gold Fiducials coverglasses (part #600-200AuF; Hestzig LLC, Leesburg, VA). A self-assembled perfusion system consisted of a DH40iL culture dish incubate system (model 640388; Warner Instruments LLC, Hamden, CT) and a quick release magnetic chamber for 25 mm low profile, round coverslips (model 641943; Warner Instruments LLC). Before each imaging session, 1 ml imager strand was pipetted in. Between each image session, samples were washed three times with buffer C (1× PBS, 500 mM NaCl, pH 8). Images were taken using a Dragonfly Spinning Disk and Super Resolution Microscope (Andor, Oxford Instruments) using a 63× oil objective and a Zyla camera at 100 ms exposure time. Image reconstruction and quantification were carried out using Picasso software following the published protocol (Schnitzbauer et al., 2017). In brief, 2000 frames of the each TIFF movie file were imported and processed under “Picasso: Localize” using Min. Net Gradient of 1000. For quantification analysis using “Picasso: Render”, we picked 30 pixel diameter circle for each cell layer, which corresponding to the size of the cell in tested aquilegia anther samples. For each sample, 10 locations were taken for each cell layer were used to calculate the number of binding sites.

## Acknowledgements

We thank members of the Meyers lab for helpful discussions, and Joanna Friesner for assistance with editing. We thank Mayumi Nakano for assistance with data handling and Parth Patel for help in the analysis of position-specific sequence variation. This work was supported by resources from the Donald Danforth Plant Science Center and the University of Missouri – Columbia. This research was supported by US NSF IOS award 1754097 to B.C.M., with additional support from funds from the Donald Danforth Plant Science Center and the University of Missouri - Columbia, plus the William H. Danforth Plant Science Fellow Award (to S.P.).

## Supplemental Figure Legends

*Supplemental Figure 1. The 24-nt phasiRNAs triggered by miR2275 in columbine* A. Above: Sequence logo of target sites of 646 24-*PHAS* loci triggered by miR2275. Below: Nucleotide sequence alignment of 24-*PHAS* loci, with sequence similarity indicated by the intensity of the blue color. Even at low resolution, it is apparent that the only target site is conserved.

B. Top: Abundance (RP10M) of small RNAs in both strands of an example locus padded with 1000 base pairs, each side. sRNA sizes are denoted by colors as indicated. Below: Phasing score of same locus and the red dot indicates the highest phased sRNA position. The cleavage site of miR2275 is denoted by the red arrow.

*Supplemental Figure 2. Aco-cand81 and miR2118/482 family members and a cluster of 24-*PHAS* loci triggered by aco_cand81*.

A. Alignment of members of the aco_cand81 family of miRNAs in columbine. The degree of conservation is indicated by the intensity of the blue color and the consensus sequence of the alignment is shown as a sequence logo

B. Same as A for miR2118 family members in columbine.

C. A cluster of ten 24-*PHAS* loci triggered by aco_cand81 in chromosome 4 spread in a 32.2 kb region. The custom browser shot shows abundances (RP10M) in top and phasing score in below tracks; the light orange region in rectangle grid shows predicted 24-*PHAS* loci region, and the red arrow indicates the cleavage site.

D. Dot plot of all of the 24-*PHAS* loci in chromosome 4 in columbine showing similarity within and between PHAS loci in nucleotide level. Green dots indicates >75% and light green dots indicates >50% similarity.

*Supplemental Figure 3. Position-specific biases in the nucleotide composition of 24-nt phasiRNAs and hc-siRNAs in columbine.*

A. Profiles of position-specific base usage of each nucleotide (A, C, G and U) comparing miR2275-derived 24-nt phasiRNAs and hc-siRNAs in columbine. All of the 24-nt phasiRNAs triggered by miR2275 with >5 raw reads (~12000 phasiRNAs) were compared with the 1000 most-abundant hc-siRNAs (matched to transposons and/or repetitive sequences). The open circle in each position indicates frequencies of each of the nucleotides, the small square boxes (highlighted with dotted circle) indicates a statistically significant (p = 1e-4) position which distinguishes two types of small RNAs.

B. Similar to A, comparing miR2118- and aco_cand81-derived 24-nt phasiRNAs with hc-siRNAs.

C. Similar to A, comparing miR2118- and aco_cand81-derived 24-nt phasiRNAs with miR2275-derived 24-nt phasiRNAs.

*Supplemental Figure 4. 24-nt phasiRNAs triggered by miR2275 in wild strawberry.*

A. Accumulation pattern of miR2275-derived 24-nt reproductive phasiRNAs in different tissues and anther development stages of wild strawberry. Athst indicates anther stage. The key at right indicates the abundance in units of log2(RP20M).

B. Alignment of members of miR2275 family in wild strawberry.

C. Abundance of miR2275 members in log10(RP20M) in different tissues of wild strawberry.

*Supplemental Figure 5. 24-nt phasiRNAs triggered by miR2275 in rose.*

A. A polycistronic precursor of miR2275 from chromosome 5 in rose. The mature and star sequences are indicated by yellow and blue respectively. The names of mature sequence were mentioned near the duplex structure, the grey highlighted names represents non-expressed candidate miR2275 sequences. The chromosomal coordinates of the precursor are indicated below the structure.

B. Alignment of miR2275 mature sequences detected in sRNA sequencing data in rose.

C. Abundance of miR2275 members in log10(RP20M) in different tissues of rose.

*Supplemental Figure 6. miR2275 in a magnoliid.*

A. Alignment of unique miR2275 mature sequences detected in genome of *Magnolia biondii*.

B. A polycistronic precursor of miR2275 from contig utg000427l in *Magnolia biondii*. The mature and star sequences are indicated by yellow and blue respectively. The names of mature sequence were mentioned near the duplex structure. The chromosomal coordinates of the precursor are indicated below the structure.

## Supplemental Tables

*Supplemental Table S1. miR2275 identified in columbine as triggers of reproductive 24-nt phasiRNAs.*

*Supplemental Table S2. miR2275, miR2118/482, and aco_cand81 triggered 24-nt reproductive phasiRNAs in columbine.*

*Supplemental Table S3. miR2275, miR2118/482, and miR11308 triggered 24-nt reproductive phasiRNAs in wild strawberry.*

*Supplemental Table S4. miR2275 triggered 24-nt reproductive phasiRNAs in rose.*

*Supplemental Table S5. miR2275 identified in Magnolia biondii and Rosa chinensis as triggers of reproductive 24-nt phasiRNAs.*

*Supplemental Table S6. Probes designed for in situ hybridization of three triggers of reproductive 24-nt phasiRNAs in Aquilegia.*

