## Supplemental Figures for "Conserved and non-conserved triggers of 24-nt reproductive phasiRNAs in eudicots"

**Supplemental Figure 1. The 24-nt phasiRNAs triggered by miR2275 in columbine**

A. Above: Sequence logo of target sites of 646 24-*PHAS* loci triggered by miR2275. Below: Nucleotide sequence alignment of 24-*PHAS* loci, with sequence similarity indicated by the intensity of the blue color. Even at low resolution, it is apparent that the only target site is conserved. B. Top: Abundance (RP10M) of small RNAs in both strands of an example locus padded with 1000 base pairs, each side. sRNA sizes are denoted by colors as indicated. Below: Phasing score of same locus and the red dot indicates the highest phased sRNA position. The cleavage site of miR2275 is denoted by the red arrow.

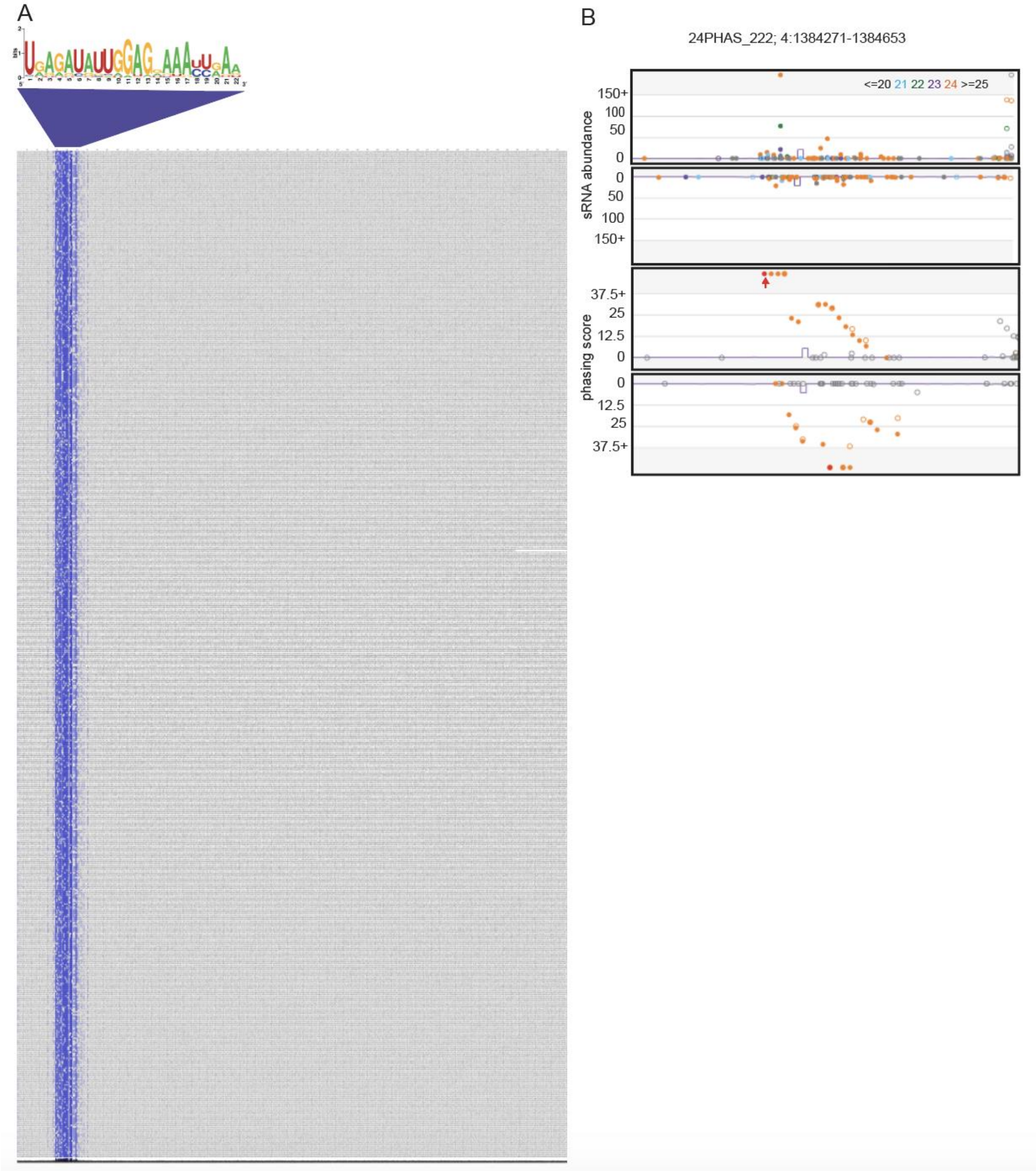

**Supplemental Figure 2. Aco-cand81 and miR2118/482 family members and a cluster of 24-*PHAS* loci triggered by aco_cand81.**

A. Alignment of members of the aco_cand81 family of miRNAs in columbine. The degree of conservation is indicated by the intensity of the blue color and the consensus sequence of the alignment is shown as a sequence logo

B. Same as A for miR2118 family members in columbine.

C. A cluster of ten 24-*PHAS* loci triggered by aco_cand81 in chromosome 4 spread in a 32.2 kb region. The custom browser shot shows abundances (RP10M) in top and phasing score in below tracks; the light orange region in rectangle grid shows predicted 24-*PHAS* loci region, and the red arrow indicates the cleavage site.

D. Dot plot of all of the 24-*PHAS* loci in chromosome 4 in columbine showing similarity within and between *PHAS* loci in nucleotide level. Green dots indicates >75% and light green dots indicates >50% similarity.

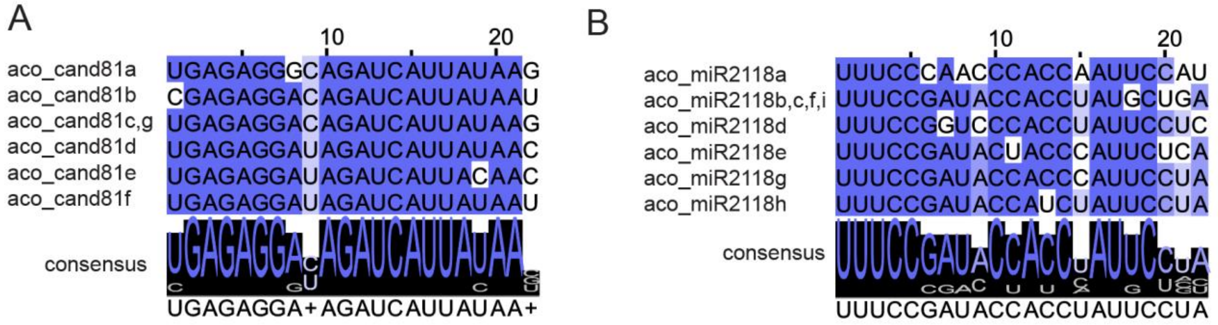

C Cluster_24PHAS; 4:2546057-2578309, 32.2 kb

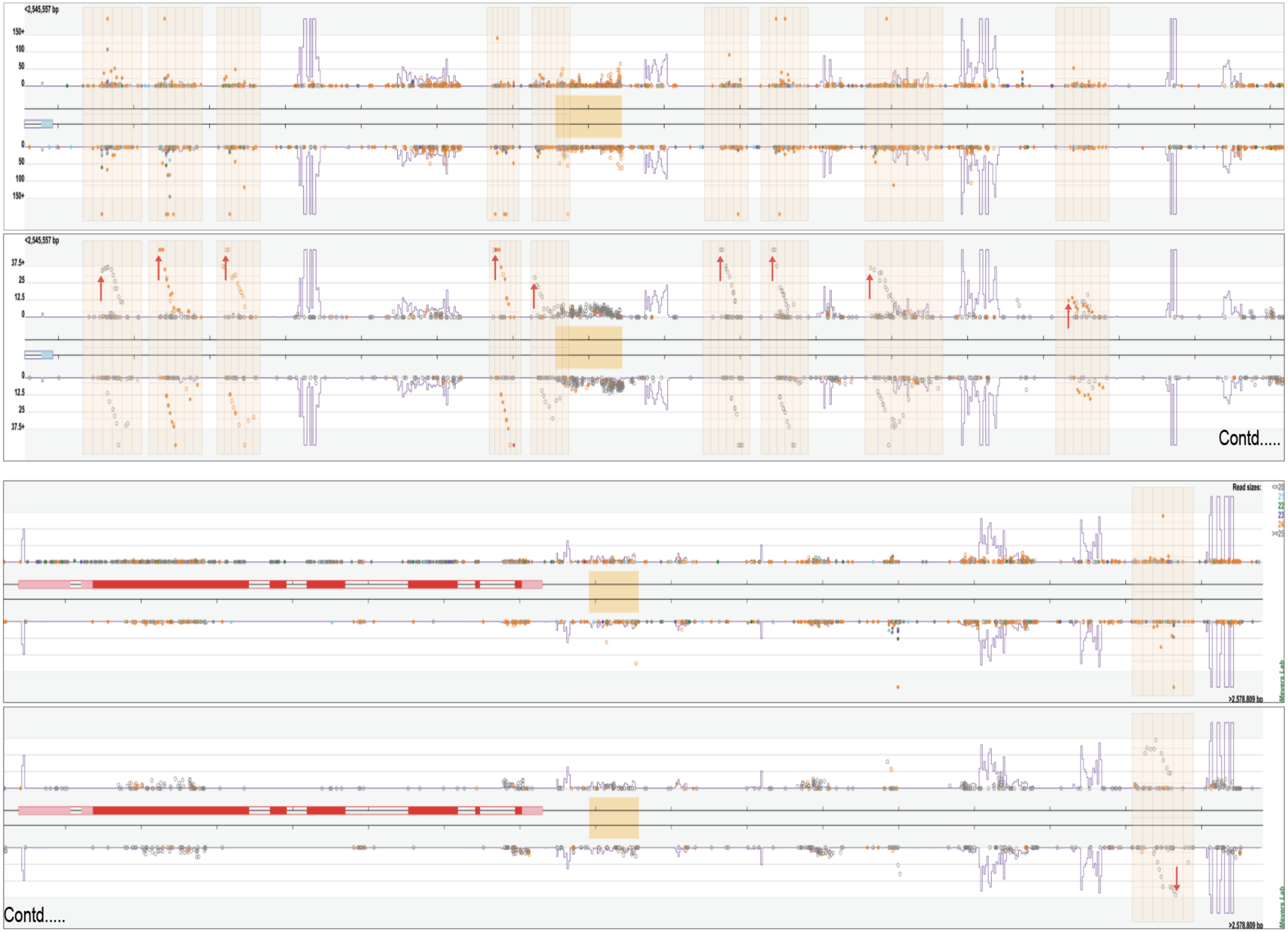

D

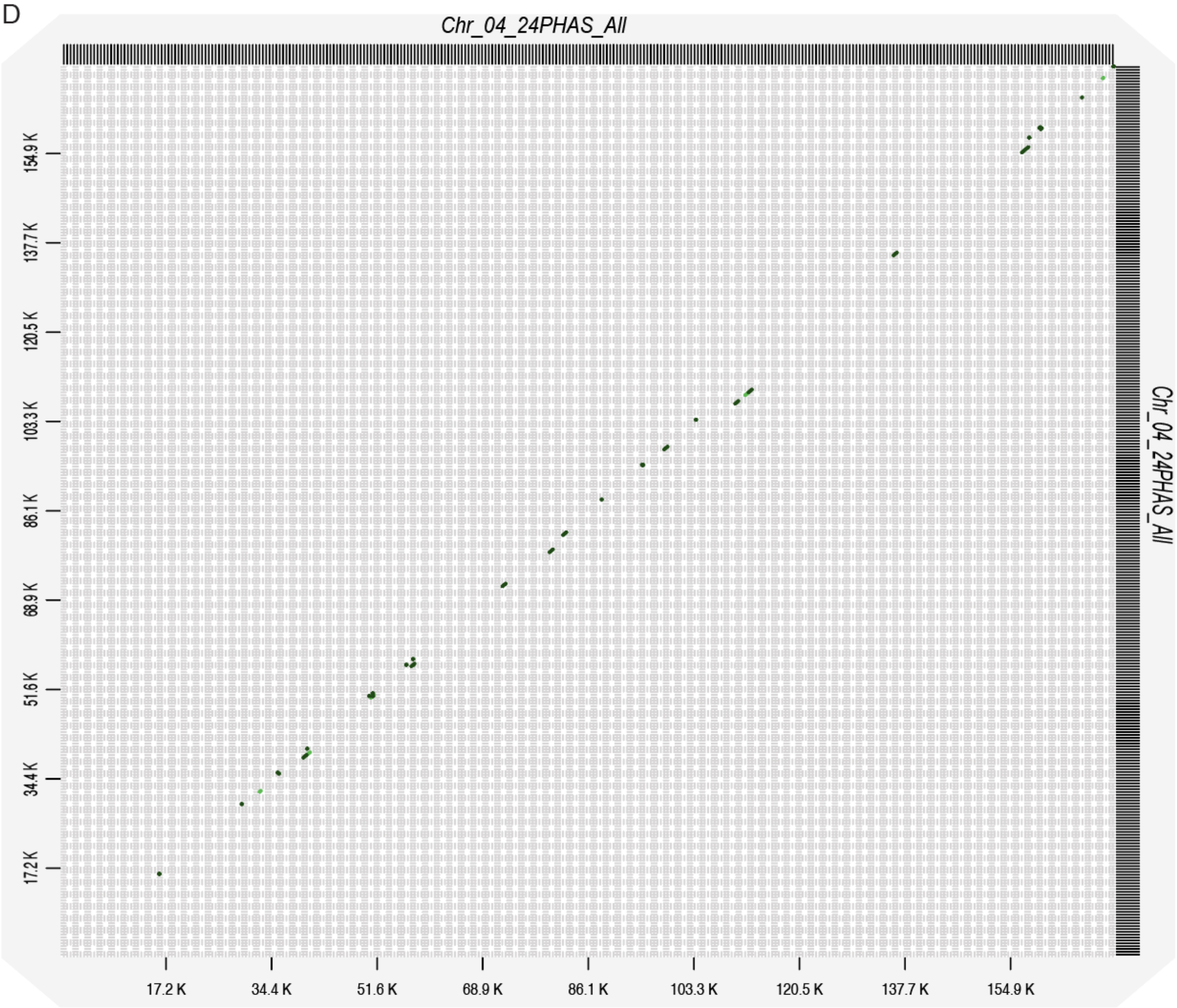

Chr_04_24PHAS_All
Chr_04_24PHAS_All
154.9 K
137.7 K
120.5 K
103.3 K
86.1 K
68.9 K
51.6 K
34.4 K
17.2 K
17.2 K
34.4 K
51.6 K
68.9 K
86.1 K
103.3 K
120.5 K
137.7 K
154.9 K

**Supplemental Figure 3. Position-specific biases in the nucleotide composition of 24-nt phasiRNAs and hc-siRNAs in columbine.**

A. Profiles of position-specific base usage of each nucleotide (A, C, G and U) comparing miR2275-derived 24-nt phasiRNAs and hc-siRNAs in columbine. All of the 24-nt phasiRNAs triggered by miR2275 with >5 raw reads (~12000 phasiRNAs) were compared with the 1000 most-abundant hc-siRNAs (matched to transposons and/or repetitive sequences). The open circle in each position indicates frequencies of each of the nucleotides, the small square boxes (highlighted with dotted circle) indicate a statistically significant (p = 1e-4) position which distinguishes two types of small RNAs.

B. Similar to A, comparing miR2118- and aco_cand81-derived 24-nt phasiRNAs with hc-siRNAs.

C. Similar to A, comparing miR2118- and aco_cand81-derived 24-nt phasiRNAs with miR2275-derived 24-nt phasiRNAs.

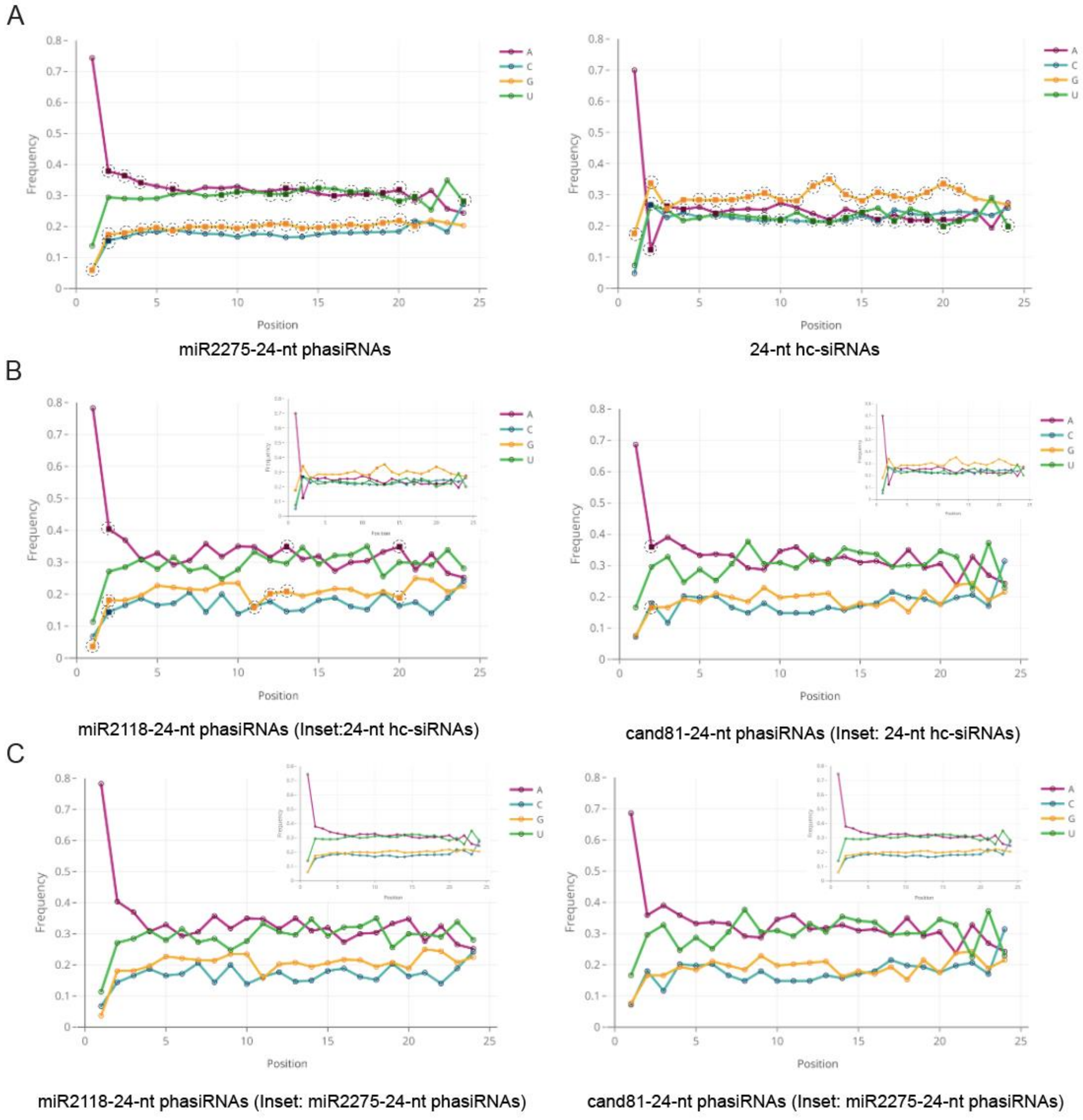

**Supplemental Figure 4. 24-nt phasiRNAs triggered by miR2275 in wild strawberry.**

A. Accumulation pattern of miR2275-derived 24-nt reproductive phasiRNAs in different tissues and anther development stages of wild strawberry. Athst indicates anther stage. The key at right indicates the abundance in units of log2(RP20M).

B. Alignment of members of miR2275 family in wild strawberry.

C. Abundance of miR2275 members in log10(RP20M) in different tissues of wild strawberry.

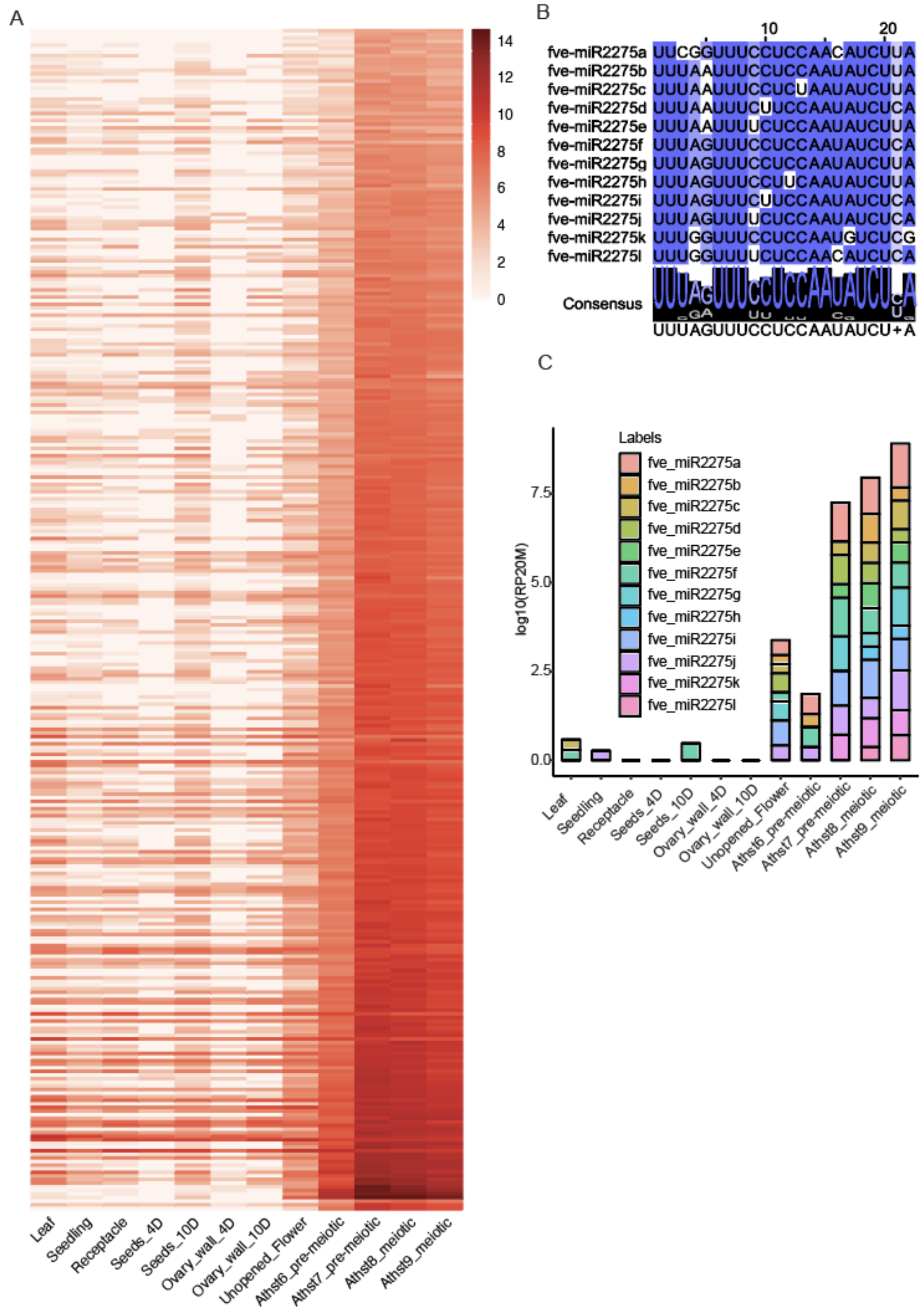

**Supplemental Figure 5. 24-nt phasiRNAs triggered by miR2275 in rose.**

A. A polycistronic precursor of miR2275 from chromosome 5 in rose. The mature and star sequences are indicated by yellow and blue respectively. The names of mature sequence were mentioned near the duplex structure, the grey highlighted names represents non-expressed candidate miR2275 sequences. The chromosomal coordinates of the precursor are indicated below the structure.

B. Alignment of miR2275 mature sequences detected in sRNA sequencing data in rose.

C. Abundance of miR2275 members in log10(RP20M) in different tissues of rose.

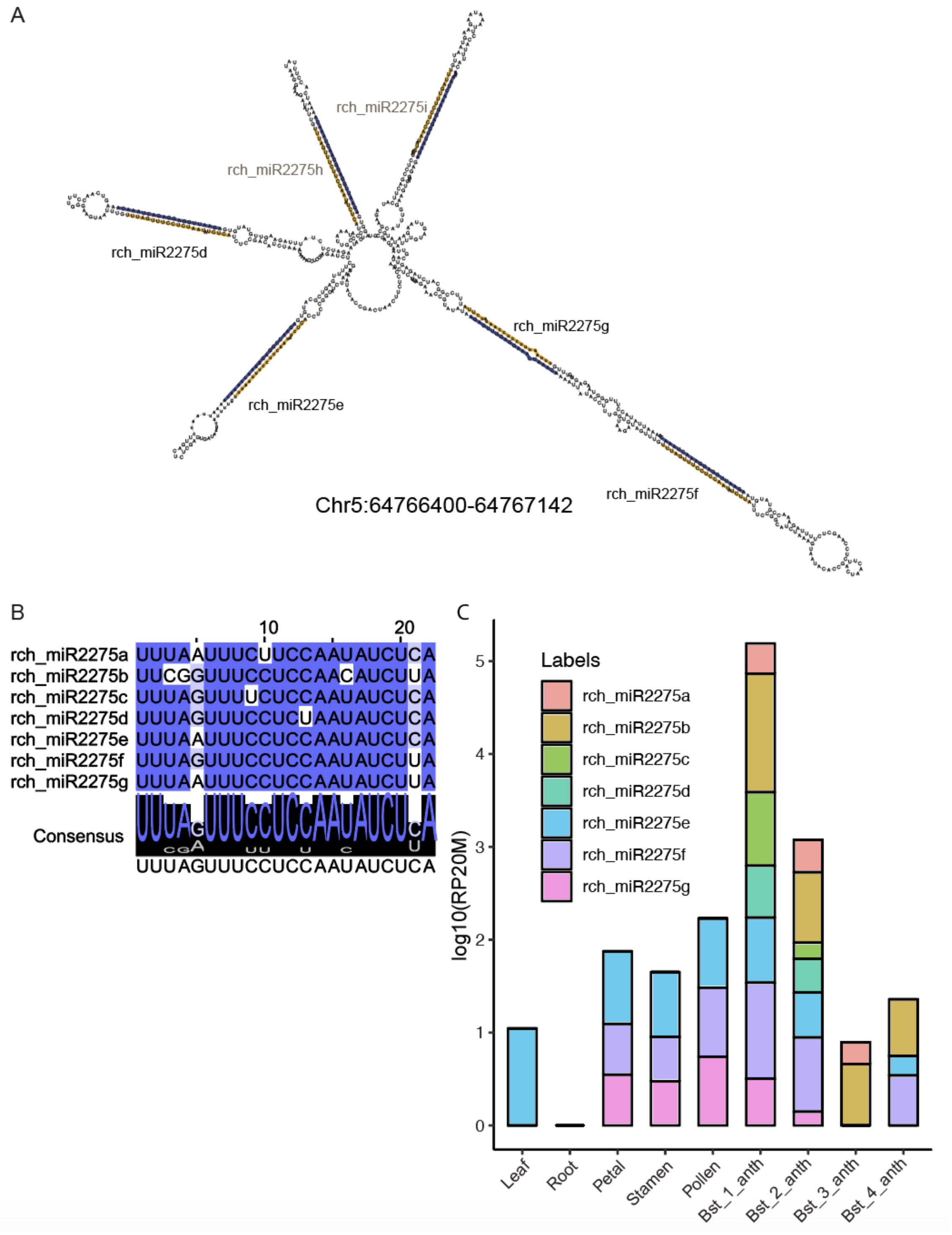

**Supplemental Figure 6. miR2275 in a magnoliid.**

A. Alignment of unique miR2275 mature sequences detected in genome of *Magnolia biondii*.

B. A polycistronic precursor of miR2275 from contig utg000427l in *Magnolia biondii*. The mature and star sequences are indicated by yellow and blue respectively. The names of mature sequence were mentioned near the duplex structure. The chromosomal coordinates of the precursor are indicated below the structure.

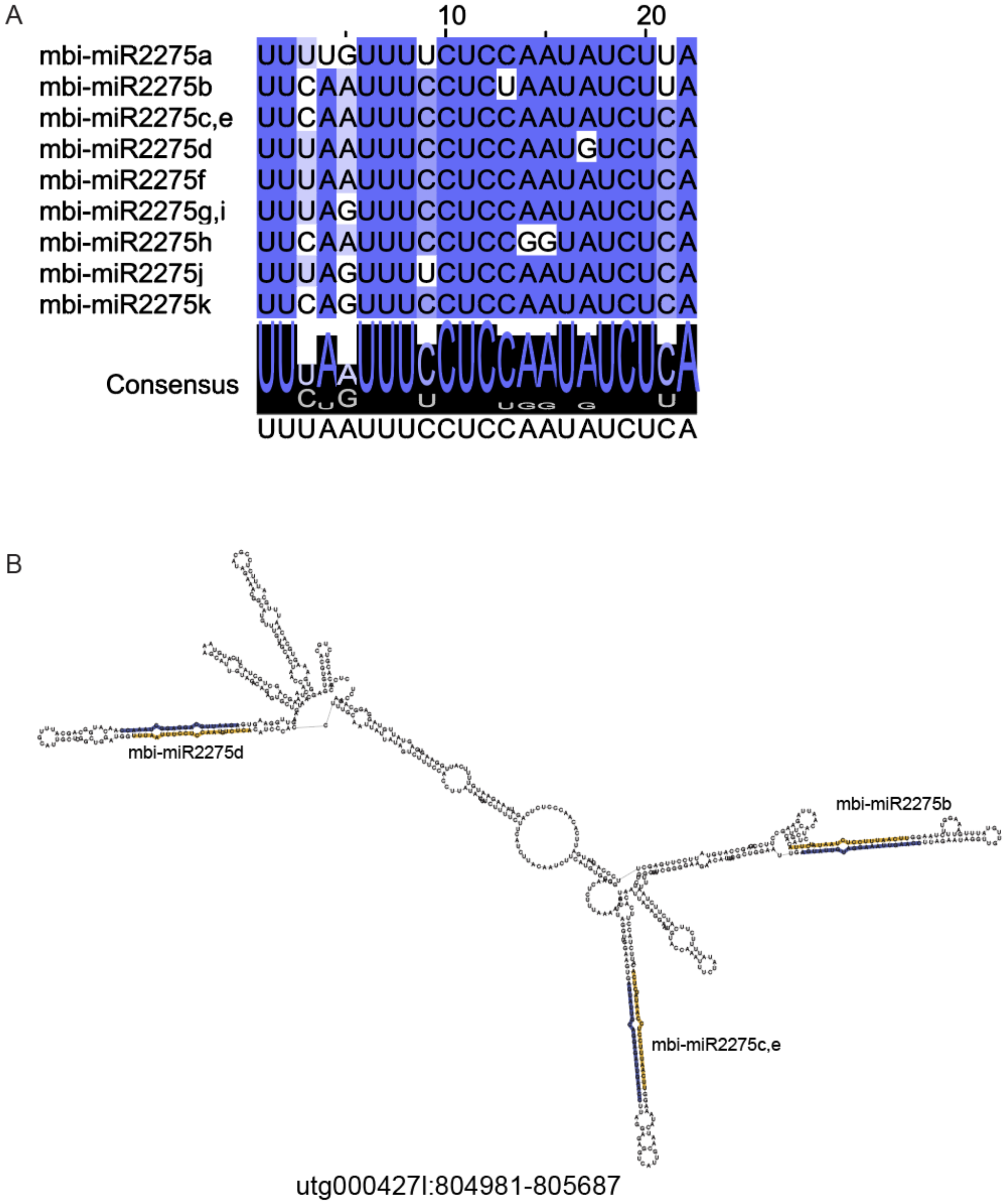
